# The E26 Transformation-Specific-family transcription factor Spi-C is dynamically regulated by external signals in B cells

**DOI:** 10.1101/2021.08.25.457658

**Authors:** Hannah L. Raczkowski, Li S. Xu, Wei Cen Wang, Rodney P. DeKoter

**Affiliations:** Department of Microbiology & Immunology and the Center for Human Immunology, Schulich School of Medicine and Dentistry, Western University, London, Ontario, Canada; Division of Genetics and Development, Children’s Health Research Institute, Lawson Health Research Institute, London, Ontario, Canada

**Author notes:** **Correspondence:** Rodney P. DeKoter, Department of Microbiology & Immunology, Schulich School of Medicine & Dentistry, Western University, London, Ontario, Canada N6A 5C1.

**Keywords:** Spi-C, transcription factor, B cell, cell fate decisions.

## Abstract

Spi-C is an E26 transformation-specific transcription factor closely related to PU.1 and Spi-B. Spi-C has lineage-instructive functions important in antibody-generating responses, B cell development, and red pulp macrophage generation. Spi-C is inducible by heme- and NF-κB-dependent pathways in macrophages. The present research aimed to examine the regulation of Spi-C expression in B cells. RT-qPCR analysis revealed that *Spic* expression was reduced in B cells following addition of lipopolysaccharide, anti-IgM antibodies, CD40L, or cytokines BAFF + IL-4 + IL-5. Cytochalasin treatment partially prevented downregulation of *Spic*. Unstimulated B cells upregulated *Spic* over time in culture. To determine the mechanism of *Spic* regulation, we examined the *Spic* promoter and upstream regulatory elements. The *Spic* promoter had unidirectional activity, which was reduced by mutation of an NF-κB binding site. *Spic* was repressed by an upstream regulatory region interacting with the heme-binding regulator Bach2. Taken together, these data indicate that Spi-C is dynamically regulated by external signals in B cells and provide insight into the mechanism of regulation.

## Introduction

At the center of adaptive immunity is the B cell. The defining phenotype of this cell subset is the expression of membrane-bound B cell receptors (BCRs) specific to antigens, and upon differentiation into plasma cells (PCs), these same immunoglobulin molecules are secreted as antibodies (Abs). In addition to the generation of Abs, activated B cells can acquire the fate of a long-lived memory B cell responsible for rapid reactivation upon secondary antigen challenge.

The E26-transformation-specific (ETS) transcription factors PU.1 (encoded by *Spi1*), Spi-B (encoded by *Spib*), and Spi-C (encoded by *Spic*) are significant contributors to cell fate decisions during hematopoiesis (1–4). PU.1 is required for generation of B cells and macrophages in mice (5–7) is and for B cells in humans (8). *Spib*^−/−^ mice exhibit far less severe defects, but show impairments in B cell development and function (1, 9). Mature B cells are not generated in mice lacking both PU.1 and Spi-B in the B cell lineage (10).

Spi-C is a lineage-instructive transcription factor that is important for the generation of multiple myeloid and lymphoid cell subsets. In the B cell compartment, Spi-C is tightly regulated during development and differentiation, functioning to promote the transition from large to small pre-B cells and regulate the generation of antibody-secreting cells (3, 11, 12). In the myeloid lineage, Spi-C is indispensable for the generation of splenic red pulp macrophages (RPMs) through a heme-dependent pathway (13, 14). Spi-C has recently been implicated in regulating the inflammatory profile of macrophages in response to NF-κB signaling, with evidence that it promotes a protective, anti-inflammatory phenotype (15). As well, altered expression of Spi-C has been noted in studies examining numerous inflammatory diseases, both in mouse models and patient cells (16, 17).

Despite its important contributions to B cell fate decisions, the mechanisms underlying the regulation of Spi-C in B cells remains largely unexplored. In macrophages, lipopolysaccharide and heme are signals that increase Spi-C expression to induce differentiation (13, 15). However, regulation of Spi-C by extracellular signals has not been assessed in B cells. In addition, there has been no work done at the molecular level to characterize regulatory elements at the *Spic* locus. The present study aimed to investigate the regulation of Spi-C in B cells. Gene expression analysis showed that *Spic* expression was reduced in B cells following addition of a variety of proliferative signals. Cultured but unstimulated B cells upregulated *Spic* over time. At the molecular level, we found that the *Spic* promoter had unidirectional activity, which was reduced by mutation of an NF-κB binding site. *Spic* was repressed by two upstream regulatory regions interacting with the heme-binding regulator Bach2. Taken together, these data indicate that Spi-C is dynamically regulated by external signals in B cells and provide insight into the mechanism of regulation.

## Results

### Heme induces Spic in primary splenic B cells and bone-marrow derived macrophages

Spi-C expression has previously been found to be induced following signaling cascades initiated by heme or NF-κB in myeloid and lymphoid cells (18). Spi-C is known to be induced in macrophages following stimulation with heme to promote differentiation into red pulp macrophages (13). We sought to determine if Spi-C expression is influenced by heme in primary B cells isolated from spleens of WT C57BL/6 mice. B cells were enriched by CD43 column depletion to 97% as determined by flow cytometry (Fig. 1A). To evaluate if *Spic* is inducible by heme in B cells, enriched B cells were cultured for 48 or 72 hours in the presence of 20 or 40 µM heme and *Spic* expression was assessed by RT-qPCR. As a control, *Spic* expression was determined in RNA prepared from freshly isolated B cells. *Spic* expression was upregulated by 3.5-fold in response to heme at 72 hours.(Fig. 1B). As prolonged culture of primary B cells is limited in the absence of pro-survival signals, we performed viable cell counts to confirm adequate cell viability. Counts indicated that the viability of B cells cultured in the presence of heme was stable over 72 hours (Fig. 1C). To validate our system of detecting *Spic* expression, we also sought to confirm upregulation of *Spic* in bone marrow-derived macrophages (BMDMs) cultured with heme reported in Haldar et al. (13). BM was isolated from WT mice and cultured for 6 days in the presence of GM-CSF, after which adherent BMDMs were plated in the presence or absence of heme (Fig. 1D). Corroborating previous findings, we found heme-treated BMDMs upregulated *Spic* expression by 5-fold compared to unstimulated cells (Fig. 1E). These results indicate that heme-mediated induction of Spi-C expression is not limited to myeloid-lineage cells and may play a role in B cell responses.

**Figure 1.**
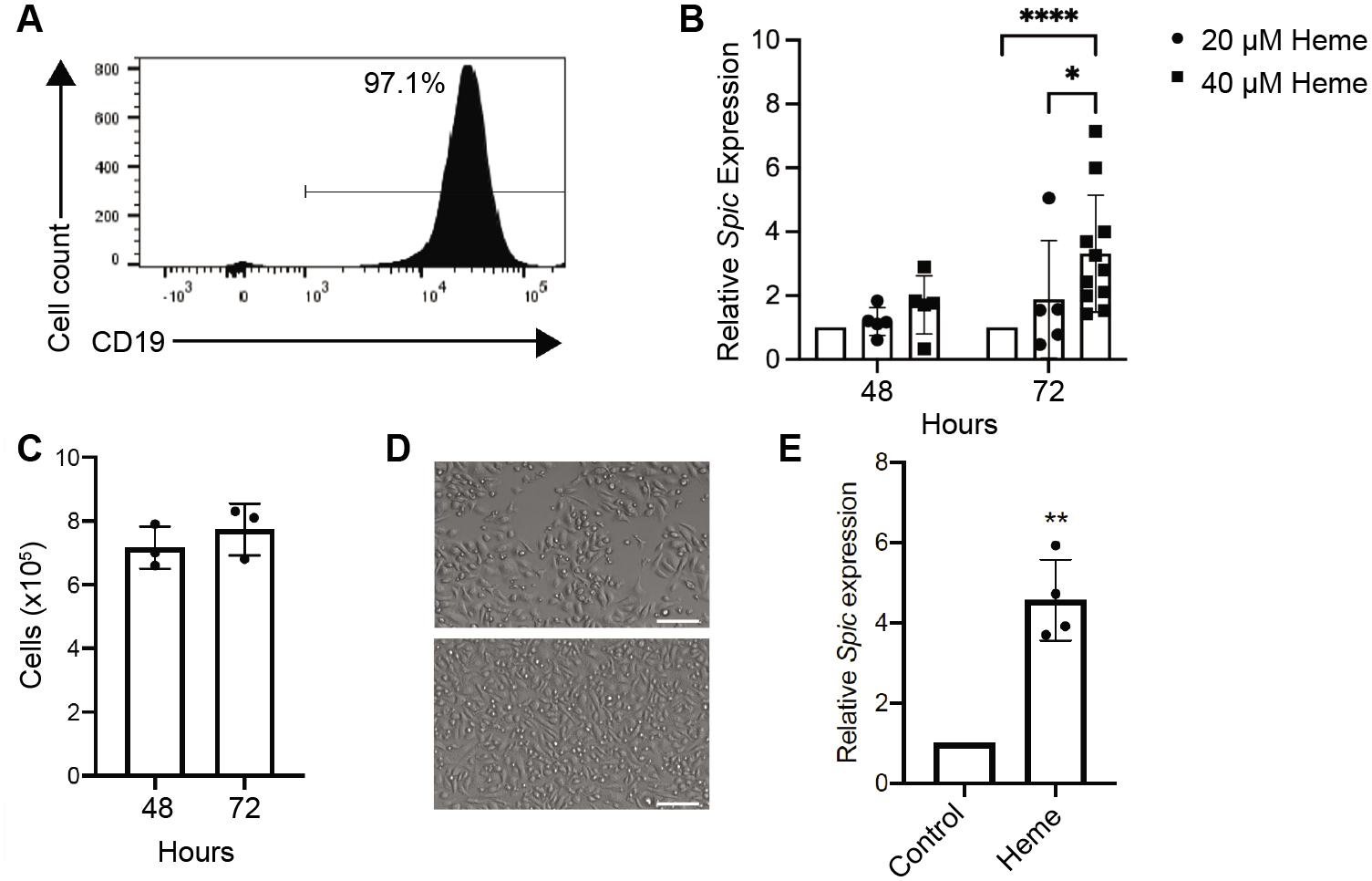
Heme induces Spic in B cells and bone-marrow derived macrophages. (**A**) Flow cytometry quantifying CD19^+^ B cell frequency following CD43-depletion of spleen cells. (**B**) RT-qPCR analysis of *Spic* expression in primary B cells enriched from WT mouse spleens and cultured with heme under the indicated conditions. Bars indicate mean ± SD. Significance was calculated by two-way ANOVA. Expression was determined relative to freshly enriched B cells. (**C**) Viable cell counts for B cells cultured in (B). Data points indicate mean of triplicates for each mouse. (**D**) Representative photomicrographs of bone-marrow macrophages cultured in cIMDM alone (top) or with 40 µM heme (bottom). Original magnification 20X. Scale bar indicates 50 µm. (**E**) RT-qPCR analysis of *Spic* expression in primary bone-marrow derived macrophages obtained from WT mice and cultured with heme for 48 hours. Bars indicate mean ± SD. Significance was determined using one sample and Wilcoxon test. Expression was determined relative to unstimulated cells cultured for the indicated amounts of time. Relative gene expression for all RT-qPCR was normalized to *Tbp*. Each data point indicates mean of duplicate wells for one mouse. \**p* < 0.05, \*\**p* < 0.01, \*\*\*\**p* < 0.0001.

### Proliferative signaling reduces Spic expression in primary splenic B cells

Various stimuli were tested to determine their impact on *Spic* expression in primary splenic B cells. Culture with IL-4 or IL-5 alone had no effect on Spi-C expression relative to cells cultured in media alone (data not shown). However, in B cells cultured with IL-4 and IL-5 in combination, a 5-fold decrease in *Spic* expression over time was observed (Fig. 2A). We noted that decreased *Spic* expression was accompanied by a slight increase in B cell numbers over time (Fig. 2B). The combination of BAFF + IL-4 + IL-5 resulted in a 40-fold decrease in *Spic* expression over 72 hours (Fig. 2C). B cells cultured in this condition also increased substantially in number over the same time course, exceeding the 10^6^ cells initially placed into culture on day zero (Fig. 2D). To examine the apparent link between B cell proliferation and *Spic* expression, we utilized the drug Cytochalasin D, a potent inhibitor of actin polymerization that therefore prevents cell division (Fig. 2D) (19). We examined *Spic* expression following culture of B cells with BAFF + IL-4 + IL-5 for 72 hours in the presence or absence of Cytochalasin D and found that addition of the drug partially blocked downregulation of *Spic* (Fig. 2E). B cells cultured with the cytokine mixture downregulated *Spic* by 16-fold, while those cultured in the presence of Cytochalasin D decreased *Spic* expression by 5-fold. This indicates that downregulation of *Spic* by BAFF + IL-4 + IL-5 is dependent on actin polymerization and blocking cell division may impair *Spic* downregulation.

**Figure 2.**
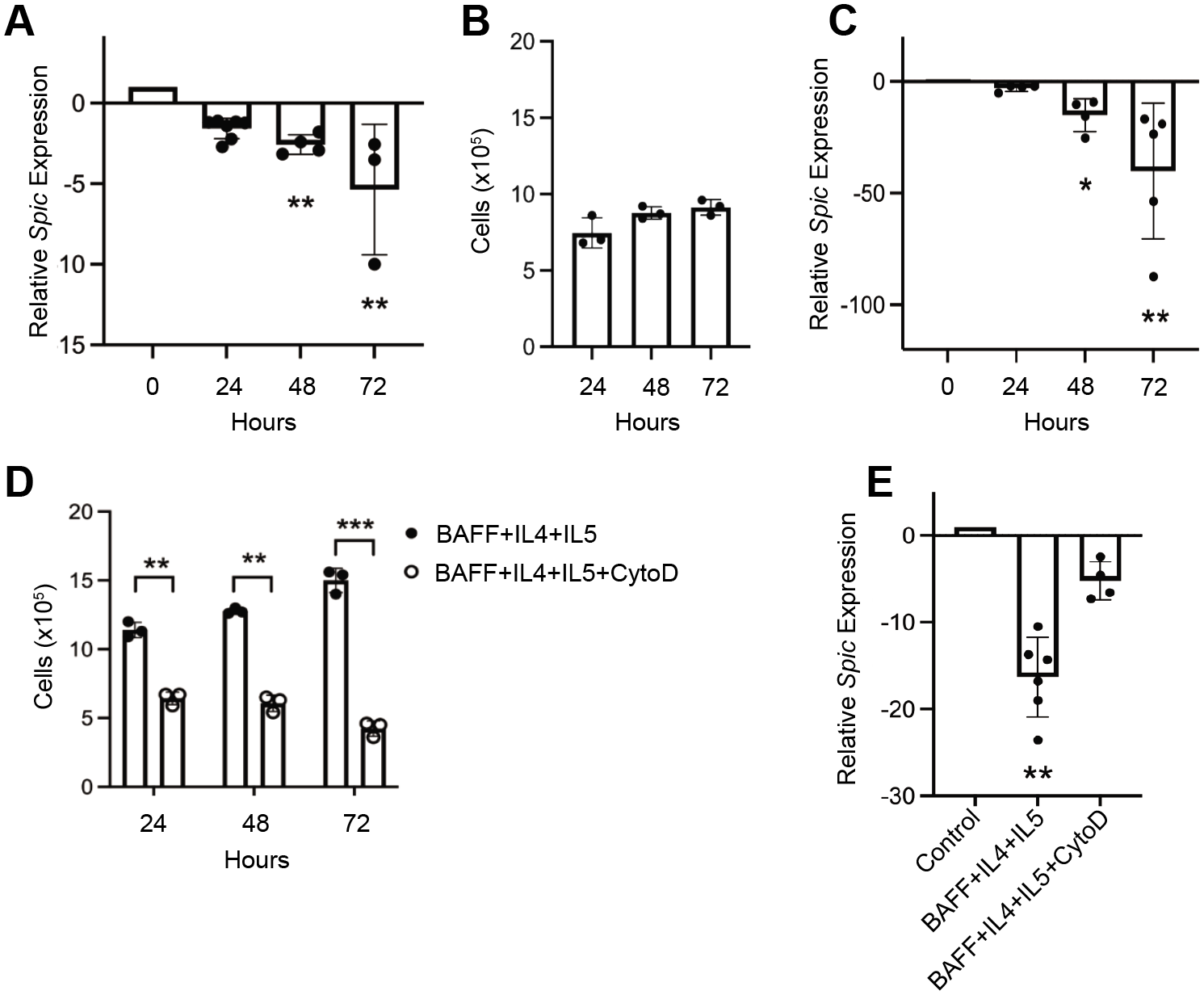
Combinations of IL-4, IL-5, and BAFF reduce *Spic* expression. (**A**) RT-qPCR analysis of *Spic* expression in primary B cells enriched from WT mouse spleens and cultured with IL-4 and IL-5. Bars indicate mean ± SD. Significance was determined by Kruskall-Wallis with Dunn’s multiple comparisons test. (**B**) Corresponding viable cell counts for (A). (**C**) RT-qPCR analysis of *Spic* expression in B cells cultured in BAFF + IL-4 + IL-5 for the indicated times. (**D**) Viable cell counts for B cells cultured in (C) or with the addition of Cytochalasin D. Bars indicate mean ± SD. Significance was determined by two-way ANOVA. (**E**) RT-qPCR analysis of *Spic* expression in B cells cultured with BAFF + IL-4 + IL-5 for 72 hours in the presence of absence of Cytochalasin D. Bars indicate mean ± SD. Significance was determined using Kruskall-Wallis with Dunn’s multiple comparisons test. Relative gene expression for all RT-qPCR was relative to freshly isolated B cells and normalized to *Tbp*. Each data point for qPCR experiments represents mean of duplicate wells for one mouse. Cell count data points indicate mean of triplicate counts for each mouse. * *p* < 0.01, \*\*\**p* < 0.001.

We previously reported that stimulation of B cell co-receptors with CD40L reduced *Spic* expression in B cells compared to freshly isolated cells (3). This observation was confirmed in the present study (Fig. 3A). We further examined the effect of CD40L by comparing *Spic* expression of stimulated cells to cultured but unstimulated B cells across three time points. *Spic* expression was reduced over time, with its lowest expression at 72-hours, showing a 29-fold reduction in expression (Fig. 3B). B cells cultured with the addition of CD40L increased in number over time (Fig. 3C). We went on to examine how CD40L affects *Spic* expression in combination with the cytokines discussed above. We found that compared to freshly isolated B cells, *Spic* was downregulated in the presence of CD40L + IL-4 + IL-5 by 16-fold compared with 7-fold using CD40L alone (Fig. 3D).

**Figure 3.**
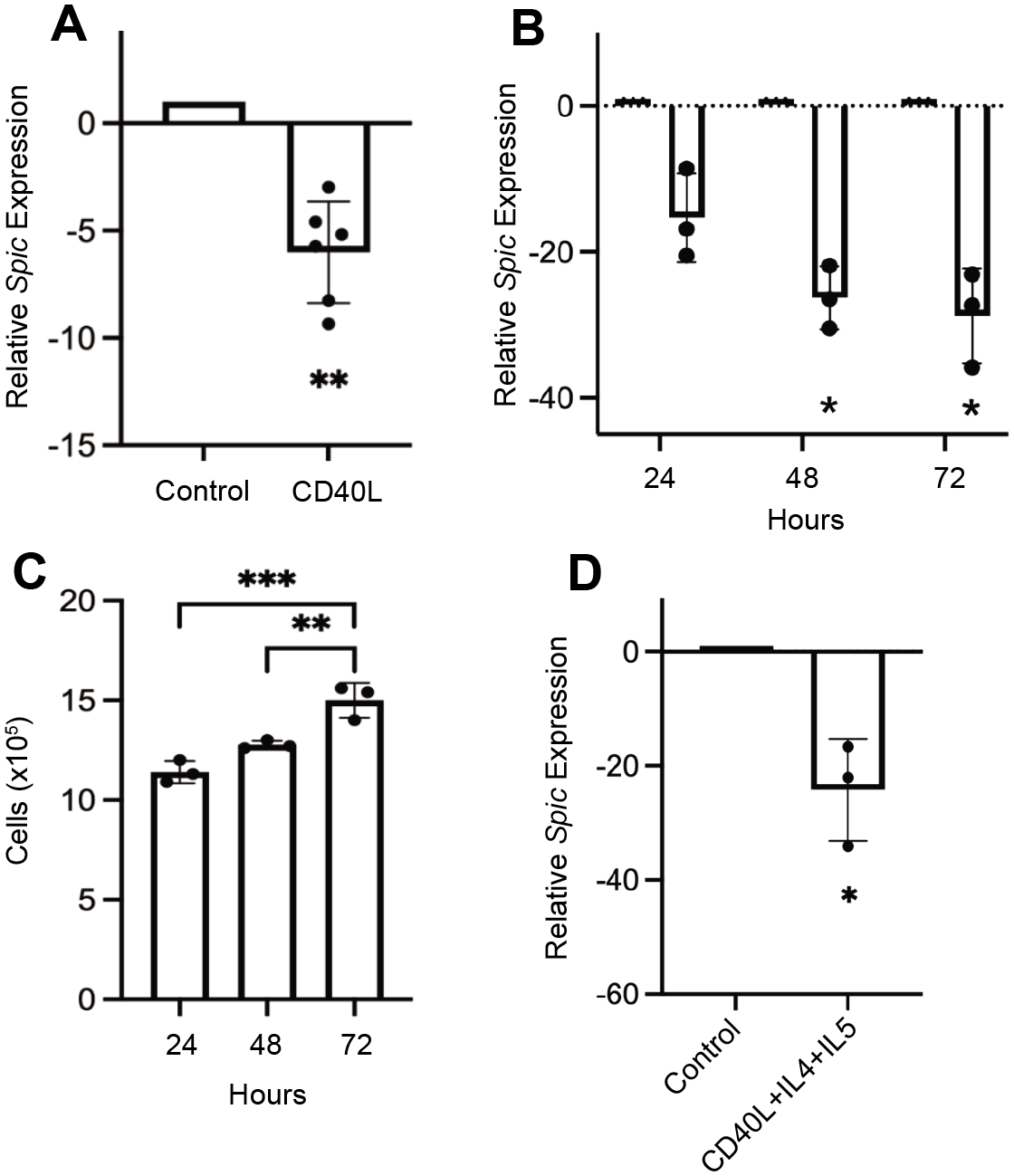
CD40L downregulates *Spic* expression in B cells. (**A**) RT-qPCR analysis of *Spic* expression in primary B cells enriched from WT mouse spleens and cultured with CD40L for 24 hours. Bars indicate mean ± SD. Significance was determined using one sample t and Wilcoxon test. (**B**) RT-qPCR analysis showing *Spic* expression in B cells cultured with or without CD40L for the indicated times. Bars indicate mean ± SD. Significance was determined using two-way ANOVA, (**C**) Viable cell counts for B cells cultured with CD40L for 72 hours. Bars indicate mean ± SD. Significance was determined using one-way ANOVA. Data points indicate mean of triplicate counts for each mouse. (**D**) RT-qPCR analysis of *Spic* expression in B cells cultured with the indicated conditions for 24 hours. Bars indicate mean ± SD. Significance was determined using one sample t and Wilcoxon test. Relative gene expression for all RT-qPCR was relative to freshly isolated B cells, with the exception of (B), which was relative to time-matched unstimulated cells. All expression data was normalized to *Tbp*. Each data point for qPCR experiments represents mean of duplicate wells for one mouse. *p < 0.05. **p < 0.01, \*\*\**p* < 0.001.

To determine the extent of the relationship between B cell proliferation and reduced *Spic* expression, we selected two additional signals to investigate their effect on *Spic* expression in B cells. We first asked how BCR signaling influenced *Spic* expression. B cells were cultured with anti-IgM Abs for 24-72 hours and *Spic* expression was quantified by RT-qPCR relative to time-matched cultured but unstimulated cells. We found that BCR engagement reduced *Spic* expression in a time-dependent manner, with expression reducing by 50-fold (Fig. 4A). As expected, stimulation through the BCR also significantly increased the number of live cells in culture over time (Fig. 4B).

**Figure 4.**
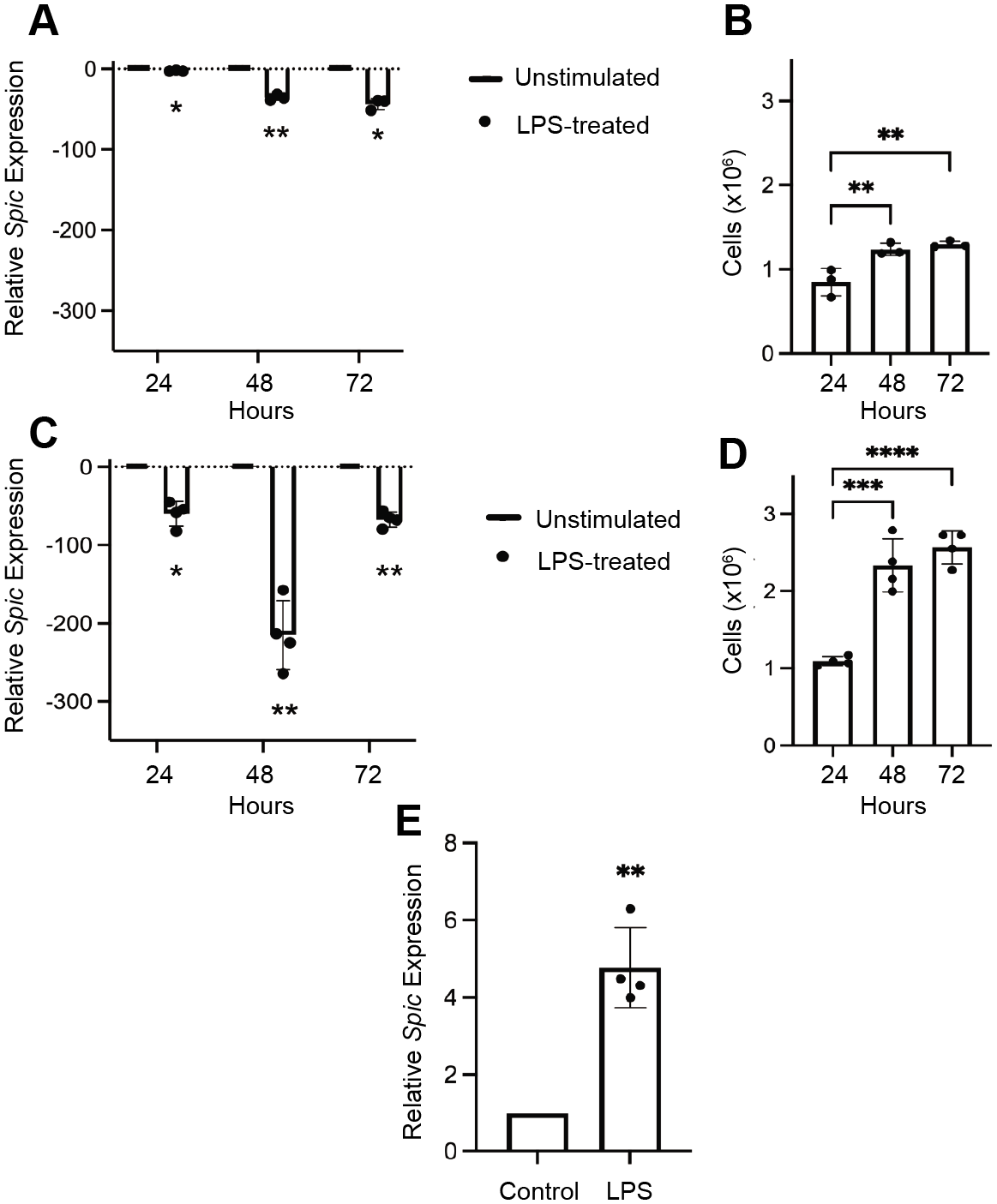
Treatment with anti-IgM Abs or LPS downregulates *Spic* expression in B cells. (**A**) RT-qPCR analysis of *Spic* expression in primary B cells enriched from WT mouse spleens and cultured with anti-IgM Abs for the indicated times. Bars indicate mean ± SD. Significance was determined using two-way ANOVA. (**B**) Corresponding viable cell counts for (A). Bars indicate mean ± SD. Significance was determined using one-way ANOVA with Tukey’s multiple comparison test. (**C**) RT-qPCR analysis showing *Spic* expression in B cells cultured with or without LPS for the indicated times. (**D**) Viable cell counts from (C). (**E**) RT-qPCR analysis of *Spic* expression in primary bone-marrow derived macrophages obtained from WT mice and cultured with LPS for 48 hours. Bars indicate mean ± SD. Significance was determined using one sample Wilcoxon test. Relative gene expression for all RT-qPCR was relative to time-matched unstimulated cells, with data was normalized to *Tbp*. Each data point for qPCR experiments represents mean of duplicate wells for one mouse. Cell count data points indicate mean of triplicate counts for each mouse. \**p* < 0.05, \*\**p* < 0.01.

Alam and colleagues recently reported that treatment of BMDMs with LPS activated *Spic* in an NF-κB-dependent pathway to push macrophages towards an anti-inflammatory phenotype and control the immune response (15). To evaluate whether a similar transcriptional program exists in B cells, we treated primary splenic B cells with LPS for 24-72 hours. We observed that LPS treatment downregulated *Spic* expression by 225-fold in B cells (Fig. 4C). *Spic* expression was lowest at 48 hours and recovered to an extent at 72 hours. Corresponding cell count data displayed a considerable increase in live B cell counts over time, peaking at over 2.5 x 10^6^ cells following 72 hours in culture (Fig. 4D). To validate our findings, we sought to reproduce the previously described reports of LPS treatment activating *Spic* expression in macrophages (Alam et al., 2020). We utilized the aforementioned culture system to generate BMDMs and plated adherent cells in the presence or absence of LPS. We found that LPS increased *Spic* expression in BMDMs by 5-fold, which confirmed the modest upregulation observed by the Haldar laboratory (15) (Fig. 4E). These data further support the correlation between B cell division and downregulation of *Spic*, including in the context of a signal that activates *Spic* expression in the myeloid lineage.

### Spic expression is increased by quiescence or anti-proliferative signaling in primary splenic B cells

Based on our previous observations, we next asked how expression of SPI-family transcription factors changes over time in unstimulated B cells. To assess *Spic* expression in cells serving as an unstimulated control, we quantified expression in cells cultured in cRPMI alone for 24-72 hours relative to freshly isolated B cells. We observed a time-dependent increase in *Spic* expression, culminating in a 20-fold increase by the 72-hour timepoint (Fig. 5A). To determine if the observed upregulation was specific to Spi-C among ETS transcription factors, we examined transcript levels of closely related family members *Spi1* and *Spib* using matched samples. Expression of *Spi1* and *Spib* increased over time, peaking at 11- and 4.3-fold increases, respectively (Fig. 5B-C). Cell count data showed stable numbers of live B cells over time after an initial decrease from 24 to 48 hours (Fig. 5D). These results suggest that *Spic* expression is increased in unstimulated B cells, relative to the related ETS transcription factors *Spi1* and *Spib*.

**Figure 5.**
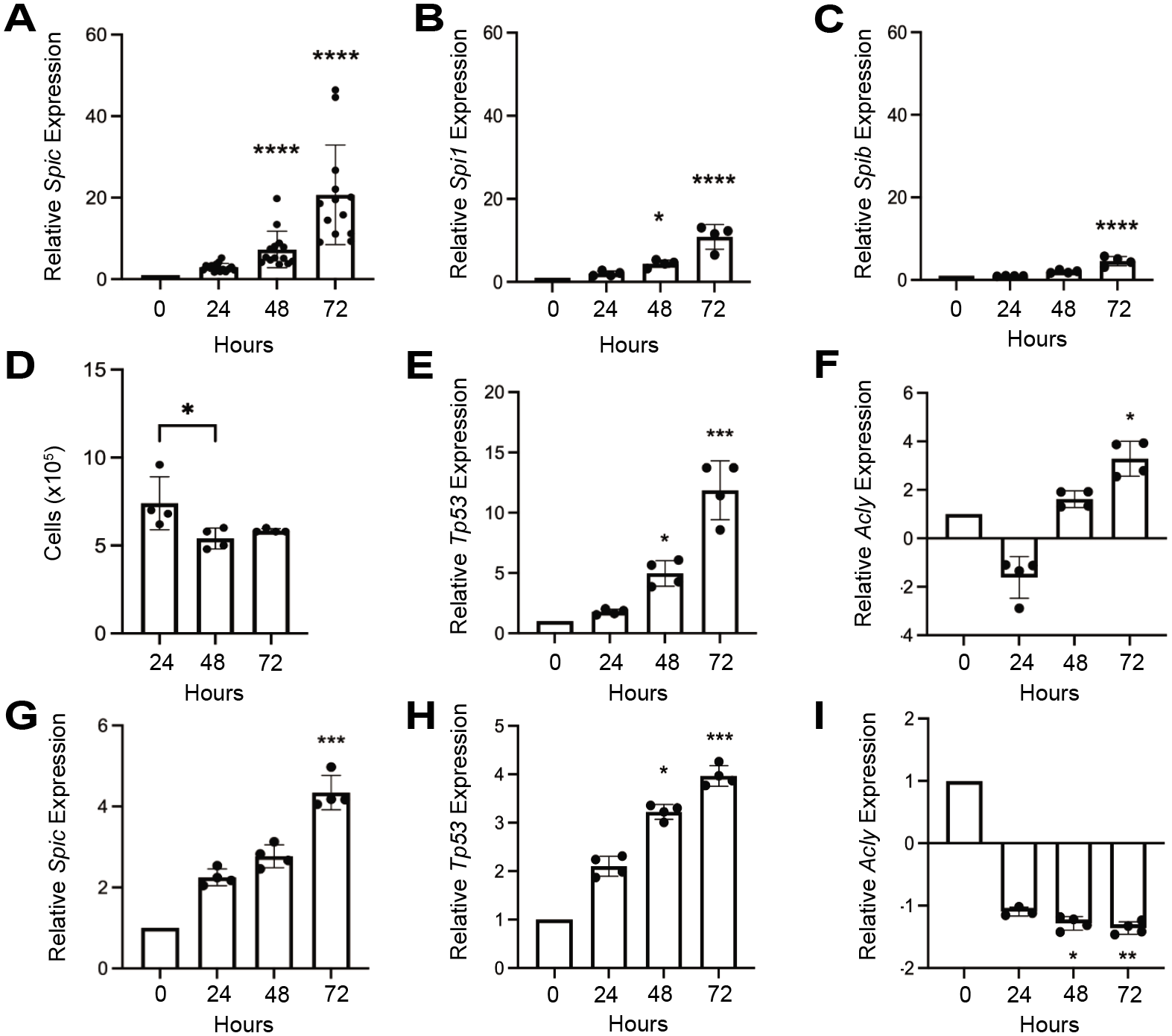
Expression of SPI family members in unstimulated B cells. (**A-C**) RT-qPCR analysis of mRNA transcript levels of *Spic*, *Spi1*, and *Spib*. WT primary splenic B cells were cultured in complete RPMI for the indicated times. Bars indicate mean ± SD. Significance was determined using Kruskall-Wallis with Dunn’s multiple comparisons test, (**D**) Live cell counts for unstimulated B cells. Bars indicate mean ± SD. Significance was determined using one-way ANOVA with Tukey’s multiple comparison test. (**E-F**) RT-qPCR analysis quantifying expression of the indicated genes in unstimulated B cells. Data shown as mean ± SD. Significance was determined using Kruskall-Wallis with Dunn’s multiple comparisons test. (**G-I**) RT-qPCR analysis quantifying expression of the indicated genes in unstimulated B cells normalized to *Actb* expression. Significance was determined using Kruskall-Wallis with Dunn’s multiple comparisons test. Relative gene expression for all RT-qPCR was relative to freshly isolated B cells. Expression was normalized to *Tbp* for all RT-qPCR, with the exception of panels G-I. Individual data points for qPCR experiments represent mean of duplicate wells for one mouse. Cell count data points indicate mean of triplicate counts for each mouse. \**p* < 0.05, \*\**p* < 0.01, \*\*\**p* < 0.001.****, *p* < 0.0001.

To further explore the upregulation of *Spic* in unstimulated B cells relative to other genes, we sought to examine the expression of genes with known patterns of expression during nutrient starvation and/or apoptosis. We selected *Tp53* as a gene that is expected to be upregulated in unstimulated B cells due to its well-documented increase in expression during apoptosis (20). *Acly* was chosen as a gene expected to be downregulated because of its role in fatty acid synthesis during cell division (21). We found that *Tp53* was upregulated over time by 12-fold in unstimulated B cells, whereas *Acly* expression decreased at 24 hours and increased by 3-fold following 72 hours in culture (Fig. 5E-F). To validate our findings, we used β-actin as an additional reference gene. We found that relative to *Actb*, *Spic* and *Tp53* transcript levels increased in a similar time-dependent manner and to the same extent, peaking at approximately 4-fold (Fig. 5G-H). Conversely, *Acly* expression decreased slightly and remained low throughout the assessed time period (Fig. 5I). Overall, these findings support the notion that *Spic* expression is increased in quiescent B cells.

Finally, we investigated *Spic* expression in B cells treated with drugs known to inhibit cell division. Primary splenic B cells were treated with Cytochalasin D, followed by quantification of *Spic* expression and cell counting. *Spic* expression increased modestly over 72 hours, with a peak increase of 3.9-fold observed at 72 hours (Fig. S1A). Cell counts remained stable over the same time period (Fig. S1B). We utilized imatinib to evaluate how inhibition of Bcr-Abl Tyrosine-kinase signaling affects *Spic* expression in B cells. Imatinib (also known as Gleevec or Glivec) is an Abl kinase inhibitor that blocks proliferation of v-Abl-transformed B cell lines (22). We treated the v-Abl-transformed pro-B cell line 38B9, and as a control treated IL-7-withdrawn fetal liver-derived WT pro-B cells, with 10 mM Imatinib for 48-72 hours and assessed *Spic* expression. Imatinib treatment induced *Spic* expression by an average of 11-fold in 38B9 pro-B cells, but not in WT pro-B cells (Fig. S1C-D). These data further support the idea of quiescence or the absence of stimulation inducing *Spic* expression in B cells.

### NF-κB activates transcription of Spic through interaction with its promoter

Next, we sought to determine the mechanism of *Spic* regulation by external signals. Spi-C expression was previously reported to be activated by NF-κB signaling in small pre-B cells (11, 12) or in macrophages (15), though no binding sites were defined. The *Spic* promoter has not been previously characterized. We selected a region immediately upstream of the transcription start site encompassing 483 bp of sequence that is highly conserved across vertebrates (Fig. 6A). This region was PCR amplified, cloned, and ligated into the pGL3-Basic luciferase reporter plasmid. CiiiDER and ConTra v3 software packages were used to identify predicted NF-κB subunit binding sites within the cloned region of the promoter (Fig. 6B-C). One common predicted NF-κB binding site was identified by both programs. We next performed site-directed mutagenesis to mutate two crucial guanine nucleotides of the RGGRNN consensus sequence known to be required for NF-κB subunit binding (23) (Fig. 6D-E). Transient transfection followed by luciferase assays in WEHI-279 cells revealed that the promoter had activity in the forward but not the reverse orientation (Fig. 6F). Transfection of the mutant vector into WEHI-279 B cells resulted in reduced luciferase activity compared to the wildtype vector (Fig. 6F). Finally, we performed an electrophoretic mobility shift assay to investigate the ability of the NF-κB protein p50 to bind to the predicted NF-κB site in the *Spic* promoter. Recombinant p50 bound to the wildtype but not the mutated NF-κB binding site, and the unmutated promoter sequence showed a supershifted complex in the presence of anti-p50 antibody. These data suggest that NF-κB activates transcription of *Spic* by binding to the identified site within the promoter.

**Figure 6.**
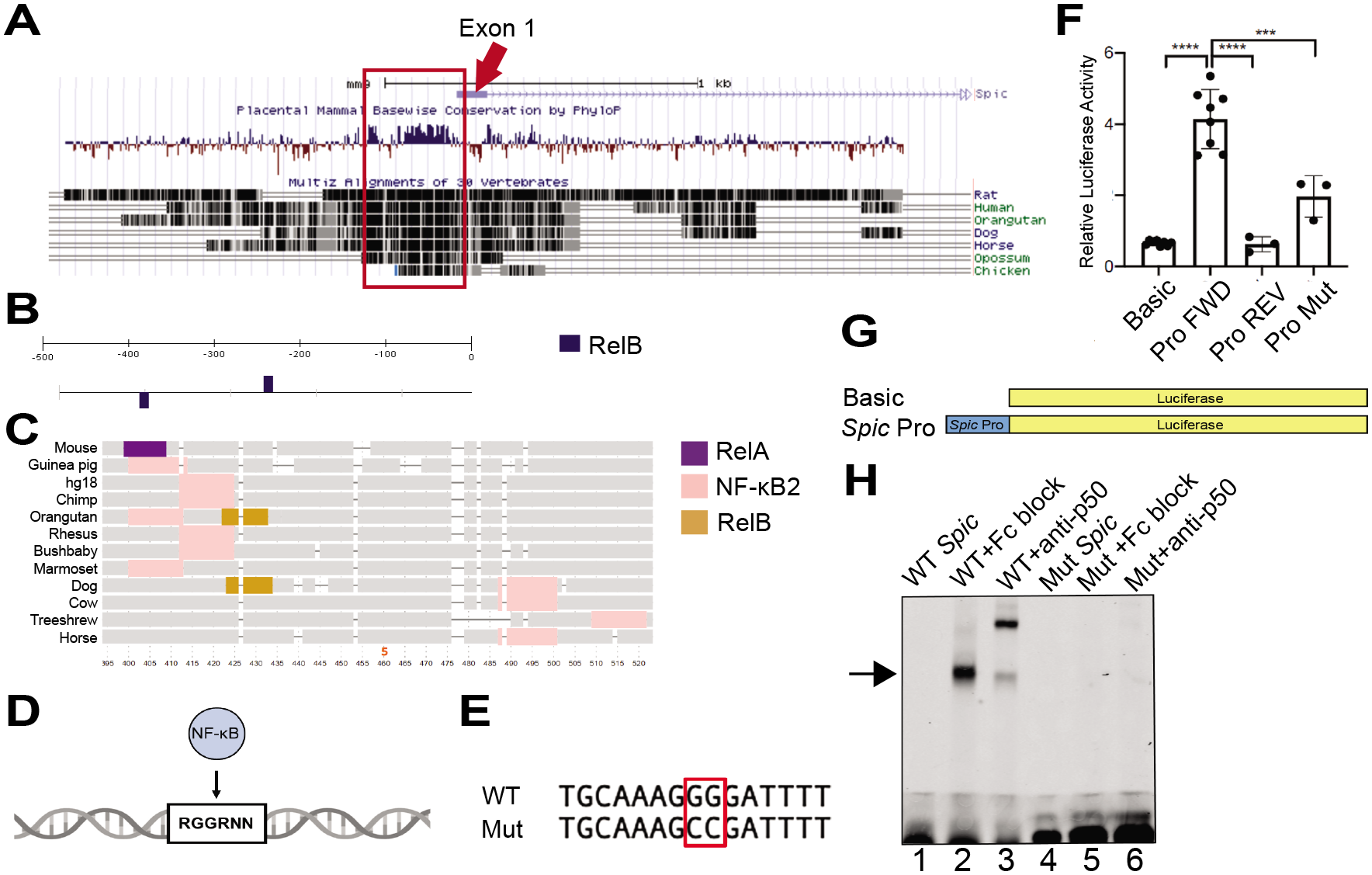
Activation of *Spic* transcription by NF-κB. (**A**) UCSC Genome Browser track showing *Spic* exon 1 and surrounding sequence. Arrow represents exon 1 and box denotes upstream region of conservation. (**B**) CiiiDER transcription factor binding site prediction within cloned promoter sequence including two possible RelB binding sites. (**C**) ConTra v3 cross-species transcription factor binding site prediction within one 110 bp region of the cloned promoter. Possible NF-κB subunit binding sites shown as coloured segments. (**D**) Schematic of NF-κB consensus binding site. (**E**) DNA sequence of cloned region of Spic promoter showing wildtype sequence and mutant. (**F**) Transient transfection of WEHI-279 B cells and luciferase assays indicated a reduction in luciferase activity following site-directed mutagenesis (top) and schematic of vectors (bottom). Relative luciferase activity (RLU) represents Renilla/Luciferase readings. Bars indicate mean ± SD. Significance was determined using one-way ANOVA with Tukey’s multiple comparisons test. Individual data points represent mean of triplicate wells for a single experiment. (**G**) Electrophoretic mobility shift assay showing binding of the WT or mutant (Mut) *Spic* promoter oligonucleotide with p50 (lane 2) and supershift in the presence of anti-p50 (lane 3). \*\*\**p* < 0.001.

### Bach2 represses transcription of Spic

Previous studies showed that *Spic* expression is repressed at the transcriptional level by Bach2 in B cells (3, 24, 25). However, Bach2 binding sites have not been identified in *Spic* regulatory regions. Reanalysis of published anti-Bach2 ChIP-Seq data (26) led to the identification of two binding peaks approximately 40 kb upstream of the *Spic* transcription start site (Fig. 7A). These two regions, termed here region of interest (ROI) 1 and 2 are in accordance with previous reports of *Spic* regulatory sequences interacting with Bach1 (25). To determine if Bach2 regulates expression of *Spic* by interacting with the identified regulatory regions, we PCR amplified ∼200 bp regions encompassing ROI 1 or ROI 2, cloned them, and ligated them into the pGL3-Spi-C promoter reporter vector (Fig. 7A). WEHI-279 B cells were transiently transfected with luciferase constructs and luciferase activity was quantified. We found that transfection of ROI 1- or 2-containing vectors alone did not enhance or repress luciferase expression relative to the vector containing only the *Spic* promoter in WEHI-279 B cells (Fig. 7B).

**Figure 7.**
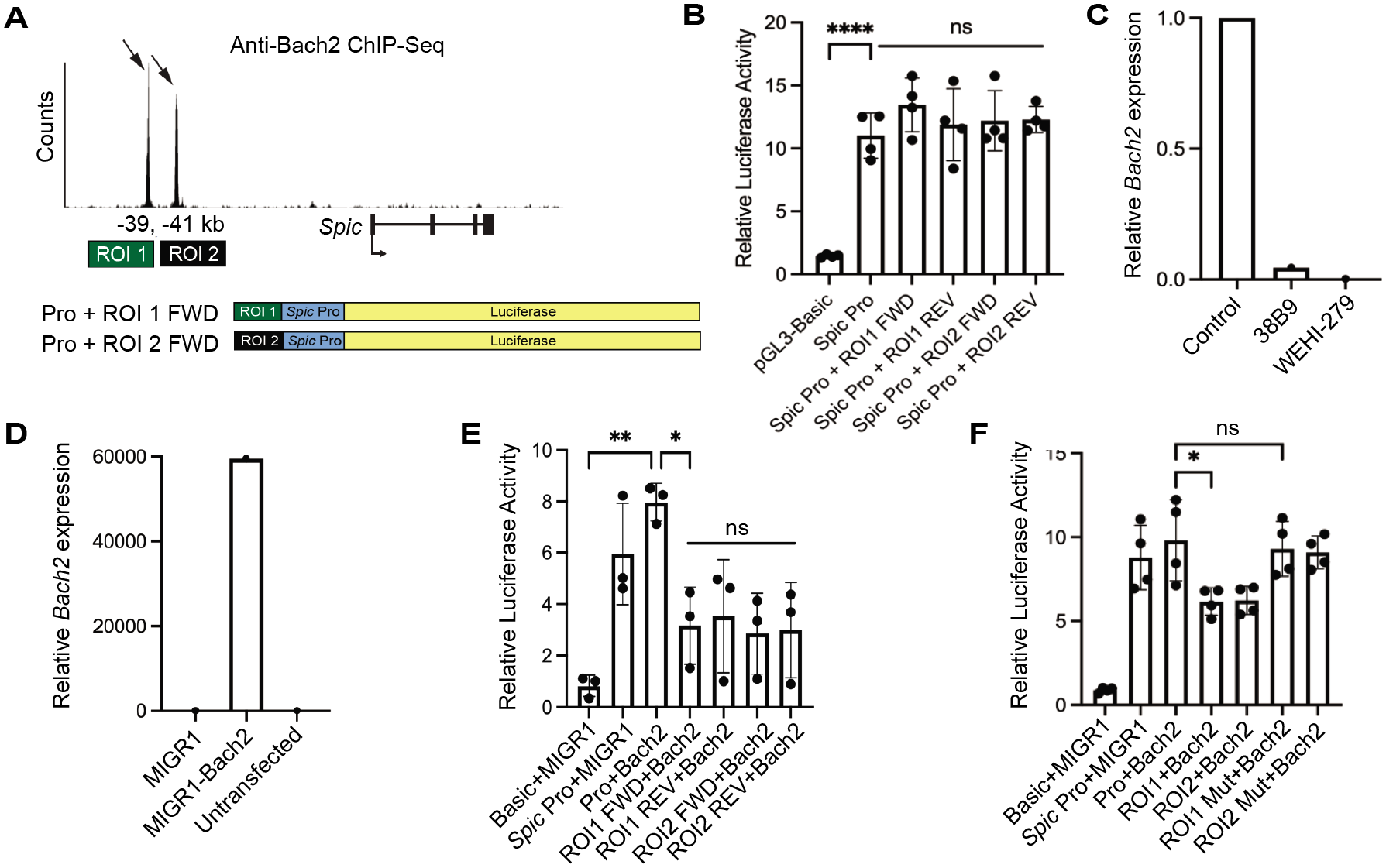
Bach2 represses *Spic* expression in B cells. (**A**) Interaction of Bach2 with regulatory regions in the *Spic* locus (top). ChIP-seq data was re-analyzed to show interaction of Bach2 with a putative regulatory element located –39 and –41 kb upstream of the Spic transcription start site. Black arrows indicate locations of Bach2 binding sites. Schematic of luciferase reporters *Spic* promoter + ROI 1 and *Spic* promoter + ROI 2 (bottom). (**B**) *Spic* ROI 1 and ROI 2 have no significant activity in WEHI-279 B cells. Relative luciferase activity represents Renilla/Luciferase readings. Significance was determined using one-way ANOVA with Tukey’s multiple comparisons test. (**C**) RT-qPCR analysis of *Bach2* expression in 38B9 pro-B cells and WEHI-279 cells compared to primary splenic B cells. Data represents a single representative experiment. (**D**) RT-qPCR analysis of *Bach2* expression in WEHI-279 B cells transfected with MIGR1, MIGR1-Bach2, or untransfected. Data represents a single representative experiment. (**E**) Relative luciferase activity of ROI 1 and ROI 2 in WEHI-279 B cells co-transfected with MIGR1-Bach2. Bars indicate mean ± SD. Significance was determined using one-way ANOVA with Tukey’s multiple comparisons test. (**F**) Relative luciferase activity of ROI 1 and ROI 2 in WEHI-279 B cells co-transfected with MIGR1-Bach2 following mutation of one Bach2 consensus binding site. Bars indicate mean ± SD. Significance was determined using one-way ANOVA with Tukey’s multiple comparisons test. \**p* < 0.05, \*\**p* < 0.01, \*\*\*\**p* < 0.0001.

We then asked whether WEHI-279 or 38B9 cell lines expressed sufficiently high levels of endogenous Bach2 to observe the effects of its interaction with the ROIs. RT-qPCR showed that both 38B9 pro-B cells and WEHI-279 mature B cells expressed low or undetectable levels of *Bach2* compared to primary splenic B cells (Fig. 7C). Therefore, we obtained a MIG-Bach2 retroviral vector allowing for co-transfection to enforce high levels of expression (Fig. 7D). Transfection of vectors containing ROI 1 or 2 in either orientation caused a significant reduction in relative luciferase activity in the presence of Bach2 (Fig. 7E). We next performed site-directed mutagenesis on one Bach2 consensus binding site within each ROI. Bach transcription factors interact with a MAF recognition element (MARE) containing the consensus sequence TGACTCA or TGAGTCA (27). A predicted MARE within ROI1 or ROI2 was mutated to replace TG with CA. Co-transfection experiments showed that vectors containing mutated Bach2 sites increased relative luciferase activity to the level observed following transfection with the *Spic* promoter (Fig. 7F). These data suggest that Bach2 represses transcription of *Spic* by interacting with ROI 1 and 2, and that this function is lost upon mutation of one Bach2 binding site.

## Discussion

The present study aimed to investigate the regulation of *Spic* by external signals and to characterize the molecular mechanisms responsible for its dynamic pattern of expression. We provided evidence that expression of Spi-C is highly sensitive to the presence of various stimuli in B cells. We showed that the absence of stimuli or use of drugs that activate a quiescent-like state upregulate *Spic*, though the exact molecular basis for this remains unknown. We also showed that NF-κB activates transcription of *Spic* by interacting with its promoter, while Bach2, a key factor involved in processes including germinal center formation and memory B cell differentiation, represses its transcription through interaction with two upstream regulatory regions. Taken together, our findings suggest that the lineage-determining transcription factor Spi-C is highly responsive to regulation by external stimuli in B cells and provide insight into the mechanism of regulation.

In general, we found that factors that induced celluar proliferation also downregulated *Spic* mRNA transcription, while factors that induced quiescence could upregulate *Spic*. CD40L (CD154) downregulated *Spic* expression in splenic murine B cells both alone and in combination with other cytokines. Compared to freshly isolated B cells, expression of *Spic* was reduced by only 5-fold in B cells cultured with only CD40L, but over 20-fold in B cells cultured with CD40L + IL-4 + IL-5. This combination of cytokines is known to simulate T cell-dependent B cell activation and induce ASC generation (28). *Spic* is known to be highly expressed in PBs and PCs; however, we previously noted that Spi-C upregulation does not occur in PB and PC differentiation induced in culture with CD40L + IL-4 + IL-5 (3, 29). Therefore, some crucial signal upregulating Spi-C expression may be missing in the culture system. It is also possible that upregulation does not occur until later in differentiation when PBs begin to lose their proliferative capacity as they mature into terminally-differentiated PCs (29).

An interesting finding was the opposing effects of LPS on *Spic* expression in B cells versus BMDMs. Of all signals assessed, LPS downregulated *Spic* expression to the greatest extent – over 200-fold – whereas LPS treatment upregulated *Spic* by approximately 5-fold in BMDMs. It is known that the NF-κB pathway becomes activated in both B cells and macrophages following TLR4 engagement (30, 31). However, one main distinction between macrophages and B cells treated with LPS is in their proliferative response. LPS-activated macrophages experience cell cycle arrest and instead respond with abundant production of pro-inflammatory cytokines and nitric oxide (30, 32). Even in the presence of M-CSF, proliferation is inhibited due to repression of cyclins (33). In contrast, B cells activated by LPS initiate a response characterized by robust proliferation and differentiation (34). Therefore, while both cell types activate the NF-κB pathway in response to LPS treatment, additional signaling events linked to the cell cycle may be responsible for differences in *Spic* expression.

We found evidence that NF-κB regulates transcription of *Spic* through a key site located in the promoter. This was suggested by work done by Alam et al. that showed that LPS signaling activates expression of *Spic* in an NF-κB-dependent manner, pushing macrophages to an anti-inflammatory phenotype to control and/or resolve the inflammatory environment (15). During B cell development, activation of the non-canonical NF-κB pathway was found to be crucial for activation of Spi-C expression during B cell development (11, 12). Previous observations from our group also identified *Nfkb1* as a target of the SPI family, with Spi-C and Spi-B repressing and enhancing its transcription, respectively (4, 9). Taken together, we propose that Spi-C exists as a key factor in a regulatory loop involving NF-κB-family members across the myeloid and lymphoid lineages, with its expressing controlling cell cycle entry during development, inflammation, and differentiation.

*Spic* was upregulated in cultured but unstimulated B cells and was upregulated to a greater extent than the related *Spi1* and *Spib* transcription factors. Previous findings show that Spi-C is highly expressed in transitional B cells, PBs, and PCs – all of which respond to BCR engagement without inducing proliferation (3, 35). Spi-C was found to directly inhibit proliferation of B cells during the pre-B cell stage of development (11, 12). Given the evidence that Spi-C is active in quiescent cells and turned off during proliferation, we propose that Spi-C is a cell cycle-responsive gene. This may be mediated in a similar fashion as seen in the regulation of RAG 1/2, which are well-established examples of cell cycle-dependent regulation (36–38). Taken together, we suggest that Spi-C directly represses cell cycle entry and is also regulated by the cell cycle to become active during periods of quiescence and repressed during proliferation.

Spi-C was induced in B cells in response to the metabolite heme. Though the molecular mechanism for its upregulation due to degradation of its repressor Bach1 is now well studied in macrophages, the biological relevance of Spi-C induction by heme in B cells remains unknown (25, 39–41). While there is evidence that treatment of B cells with heme increases transcription of the endosomal transporter HRG-1, the mechanism of heme entry into the plasma membrane is not known (40). Despite early studies reporting that phagocytosis and pinocytosis occur rarely in lymphocytes, there is growing evidence that B cells may have a higher capacity for non-specific uptake than previously thought (42, 43). Therefore, it is plausible that B cells pinocytose free heme. Alternatively, heme may be sensed externally as a danger-associated molecular pattern that drives a Spi-C-mediated immune response (3, 44). While not thought to strongly activate PRRs, there is evidence that heme may be able to signal weakly through TLRs such as TLR4 (45). Finally, it is also possible that B cells synthesize intracellular heme. These possible mechanisms of detection of heme by B cells provide insight into how it may be sensed, but further investigation of the downstream signaling pathways is warranted.

Since free heme is a potent catalyst of reactive oxygen species generation, it is maintained almost exclusively complexed with hemoglobin (44, 46). High levels of free heme are typically indicative of excessive hemolysis, which can arise due to disease or infection. It is conceivable that heme-dependent activation of Spi-C is one signaling pathway that promotes the generation of ASCs in response to a nonspecific threat (40). Specifically, we propose that in the case of hemolytic infections, such as malaria or some forms of *Streptococcus*, B cells detect and become activated in response to free heme (44, 47). This leads to the proteosome-dependent degradation of Bach2, which frees *Spic* from constitutive repression (3). Spi-C may then act to counter its related factor Spi-B and promote B cell differentiation into ASCs. In addition, we previously showed that Spi-C represses Bach2 transcription (3). Therefore, mutual cross-antagonism between Spi-C and Bach2 is a potential mechanism to promote the rapid generation of antibodies to initiate immunity while the longer-term immune response begins to develop.

In summary, this work characterized the regulation of the lineage-instructive transcription factor Spi-C in response to external signals in B cells. While the downstream effects of Spi-C in B cell development and differentiation have been described to some extent, this is to our knowledge the first study to examine its upstream regulation in mature B cells. Understanding how Spi-C expression is regulated by external signals and downstream signaling pathways in B cells will enhance knowledge of adaptive immune responses.

## Experimental Procedures

### Mice

C57BL/6 mice were purchased from Charles River Laboratories (Pointe-Claire, QC, Canada). All animals were housed under specific pathogen-free conditions at the West Valley facility (London, ON), and were monitored in accordance with an animal use protocol approved by the Western University Council on Animal Care.

### B Cell Enrichment

Spleens were removed from male and female mice aged 6-12 weeks and dissociated into a single cell suspension with ground glass tissue homogenizers. Red blood cells were lysed with ammonium-chloride-potassium buffer and B cells were enriched by negative selection using the Miltenyi system comprised of the QuadroMACS™ Separator magnet, LD depletion columns, streptavidin microbeads (Miltenyi Biotec, Germany) and biotin-conjugated mouse anti-CD43 (BD Biosciences; clone S7). Effective enrichment was confirmed by flow cytometry with staining for CD19.

### Cell Culture

Primary mouse B cells were cultured in RPMI-1640 (Wisent, St-Bruno, QC) containing 10% fetal bovine serum, 10 units penicillin/1 mg/mL streptomycin/20 mM L-glutamine, and 10^-5^ M β-mercaptoethanol (βME). Additional reagents used for culture of primary B cells are listed in Supplemental Table 1. Bone marrow was flushed from the femur and tibia of C57BL/6 WT mice aged 6-10 weeks. Following erythrocyte lysis, BM cells were plated at 2 x 10^5^ in 6-well plates and cultured for 6 days in IMDM + 10% fetal bovine serum (Wisent) supplemented with 20 ng/mL of GM-CSF (Peprotech, Rocky Hill, NJ). BMDMs were washed twice with D-PBS (Wisent) to remove non-adherent cells and cultured in fresh IMDM + 10% FBS alone or containing 1000 ng/mL LPS (List Biological Laboratories, Campbell, CA) or 40 uM hemin (Sigma-Aldrich, St. Louis, MO). After 48 hours, BMDMs were harvested for RNA extraction. WEHI-279 B lymphoma cells were cultured in DMEM containing 4.5 g/L glucose, 10% fetal bovine serum, 10 units penicillin, 1 mg/mL streptomycin, 20 mM L-glutamine, and 10^-5^ M βME. 38B9 pre-B cells were cultured in RPMI-1640 containing the same supplements as for WEHI-279. All cells were cultured at 37°C and 5% CO_2_. Images were taken with the Zeiss AxioObserver and A1 AxioCam ICM1 using ZEN 2 Pro software.

### Plasmids and Cloning

The *Spic* promoter, region of interest (ROI) 1, and ROI 2 were amplified from C57BL/6 genomic DNA by PCR using the Q5® High-Fidelity DNA Polymerase (New England BioLabs, Ipswich, MA). PCR products were ligated into pSCB-Amp/Kan using the StrataClone Blunt PCR Cloning Kit (Agilent Technologies, La Jolla, CA). The *Spic* promoter was cloned into pGL3-Basic (Promega, Madison, WI) using HindIII cut sites. Predicted NF-κB subunit binding sites within the *Spic* promoter were identified using CiiiDER (48) and ConTra v3 (49) software. Site-directed mutagenesis was performed on one common predicted site. *Spic* ROI 1 and ROI 2 were each ligated into the *Spic* promoter-containing pGL3-Basic vector using KpnI/SacI and XhoI/SacI sites. Site-directed mutagenesis was performed on one predicted Bach2 binding site in each construct. Predicted Spi-C binding sites were mutated by site-directed mutagenesis. Ligations were performed with T4 DNA Ligase (New England BioLabs). All PCR products were purified with the QIAEX II Gel Extraction Kit (Qiagen, Hilden, Germany) prior to subsequent cloning. Site-directed mutagenesis was performed using the Q5® Site-Directed Mutagenesis Kit (New England BioLabs). Constructs were verified by Sanger DNA Sequencing at the London Regional Genomics Center. Each cloned region was cloned and investigated in both the forward and reverse orientations. All restriction enzymes were purchased from New England Biolabs. Cloning and mutagenesis primers are listed in Supplemental Table 2.

### Transient Transfection

WEHI-279 or 38B9 B cells in early log-phase growth were washed three times in serum-free DMEM (4.5g/L glucose) or RPMI-1640 (Wisent, St-Bruno, QC). Cells were incubated for 10 minutes at room temperature with 0.35 µg of pRL-TK (Promega) and either 10 µg of each luciferase reporter vector or 5 µg of each reporter and 5 µg of an additional expression vector. Samples were electroporated at 220 V and 950 mF in 4-mm gap cuvettes (ThermoFisher Scientific, Rochester, NY) using a GenePulser II with Capacitance Extender (Bio-Rad). Cells were recovered at room temperature for 10 minutes and plated in 6-well culture plates in complete DMEM or RPMI for 24 hours at 37°C, 5% CO_2_.

### Luciferase Assays

Cells were washed twice in D-PBS (Wisent) and lysates were collected using the Dual-Luciferase Reporter Assay System (Promega) according to the manufacturer’s instructions. Luminescence was measured in 96-well opaque, white plates using a Synergy H4 plate reader (BioTek, Winooski, VT). Data was collected using Gen5 software (BioTek).

### Reverse transcription-quantitative PCR

Total RNA was extracted from fresh or cultured primary B cells using the RNeasy Minikit (Qiagen) or Trizol reagent (Ambion, Austin, TX). cDNA synthesis (iScript cDNA Synthesis Kit, Bio-Rad, Mississauga, ON) was performed using equal starting RNA concentrations, followed by RT-qPCR analysis, which was conducted using the SensiFAST SYBR No-ROX Kit (Bioline, Singapore) on the QuantStudio 5 or QuantStudio 3 Real-Time PCR System (ThermoFisher Scientific). Relative transcript levels were normalized to TATA-binding protein (*Tbp*) and/or β-actin (*Actb*) and calculated as fold change using the comparative threshold cycle (2-ΔΔCT) method. Primer sequences are listed in Supplemental Table 3.

### Electrophoretic Mobility Shift Assay

Forward and reverse fully complementary oligonucleotides were synthesized with 5’ conjugation to IR700 dye to contain the predicted NF-κB site in the *Spic* promoter (GCTGCAAAG**GGGATTTT**TTTT, where bold indicated the consensus binding site) or a mutant predicted to be unable to bind NF-κB (GCTGCAAAG**<u>CC</u>GATTTTT**TTT, where underlined nucleotides represent changes). Recombinant GST-p50 subunit D434-969 was purchased from Sigma-Aldrich. Binding reactions were performed for 20 minutes at room temperature with 20 pmol annealed primers. Binding buffer contained 10 mM Tris-Cl pH 7.5, 50 mM NaCl, 1 mM DTT, 1 mM EDTA, 1% Ficol-400, 1 μg poly-dI-dC (LightShift, Thermo-Fisher Scientific) in the presence of 1 μl either control antibody (2.4G2, BD Biosciences) or anti-NF-kB p50 antibody (4D1, Biolegend). Protein-DNA complexes were run on 4% non-denaturing gels for one hour using TGE running buffer and visualized using a Li-Cor Odyssey system at 700 nm.

### Statistical Analyses

All statistical analyses were performed using Prism 9.1.2 (GraphPad, La Jolla, CA). Statistical tests used are indicated in the figure legends. All figures show individual data points.

## Supporting information

This article contains supporting information (one supplementary figure and three supplemental tables).

## Author contributions

H. L. R. conceptualized the study, performed experiments, and wrote the manuscript. L. S. X. and W. C. W. performed experiments. R. P. D. conceptualized the study, edited the manuscript, and acquired funding.

## Funding and additional information

This work was supported by Natural Sciences and Engineering Research Council grant 04749-2010.

## Conflict of interest

The authors declare no conflicts of interest.

## Abbreviations

Ab: antibody
ASC: antibody-secreting cell
Bach: BTB and CNC homology
BCR: B cell receptor
BCLAF: Bcl2-associated factor
BM: bone marrow
BMDM: bone marrow-derived macrophage
cDNA: complementary deoxyribonucleic acid
ETS: E26-transformation-specific
FO: follicular
GC: germinal center
IL: interleukin
IRF: interferon regulatory factor
LPS: lipopolysaccharide
MARE: MAF recognition element
mRNA: messenger ribonucleic acid
PRR: pattern recognition receptor
PB: plasmablast
PC: plasma cell
RPM: red pulp macrophage
TLR: toll-like receptor
WT: wildtype

---

**Supplemental Table 1.**
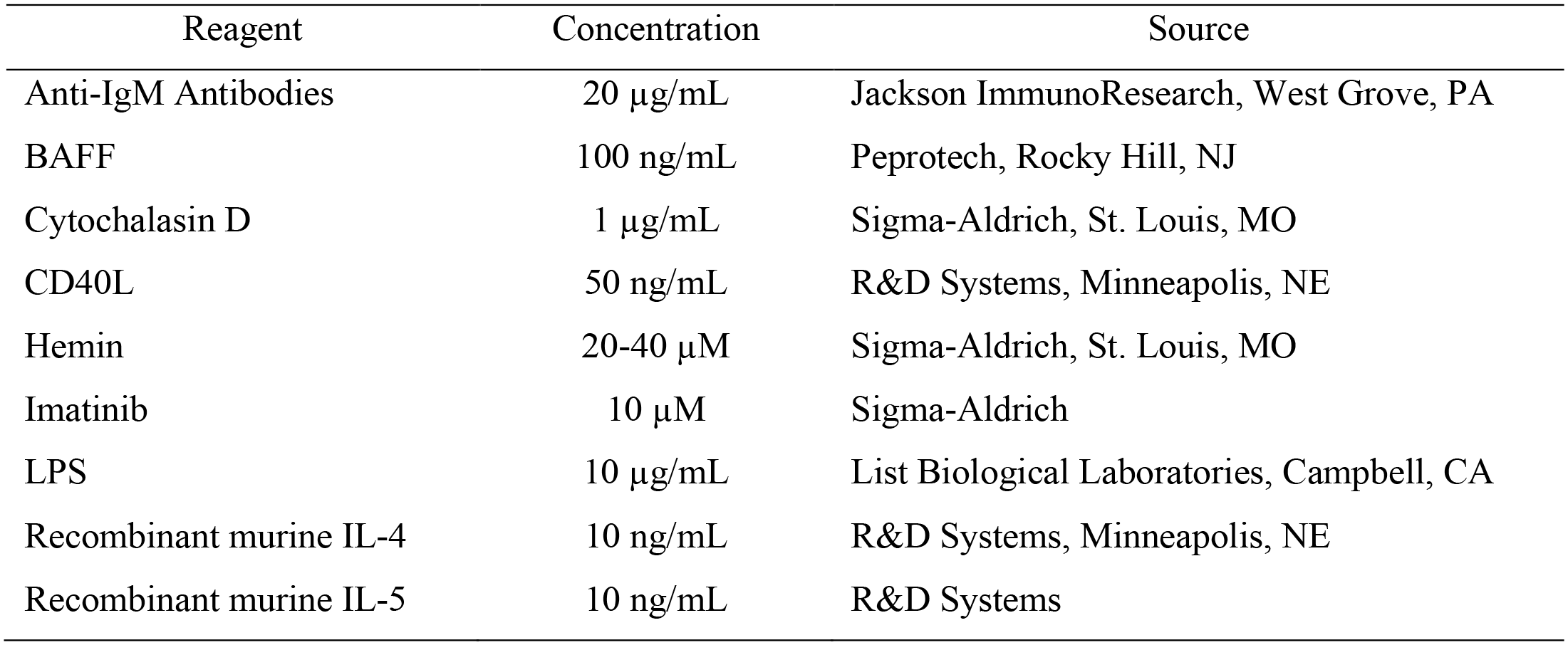
Reagents used in primary B cell culture.

**Supplemental Table 2.**
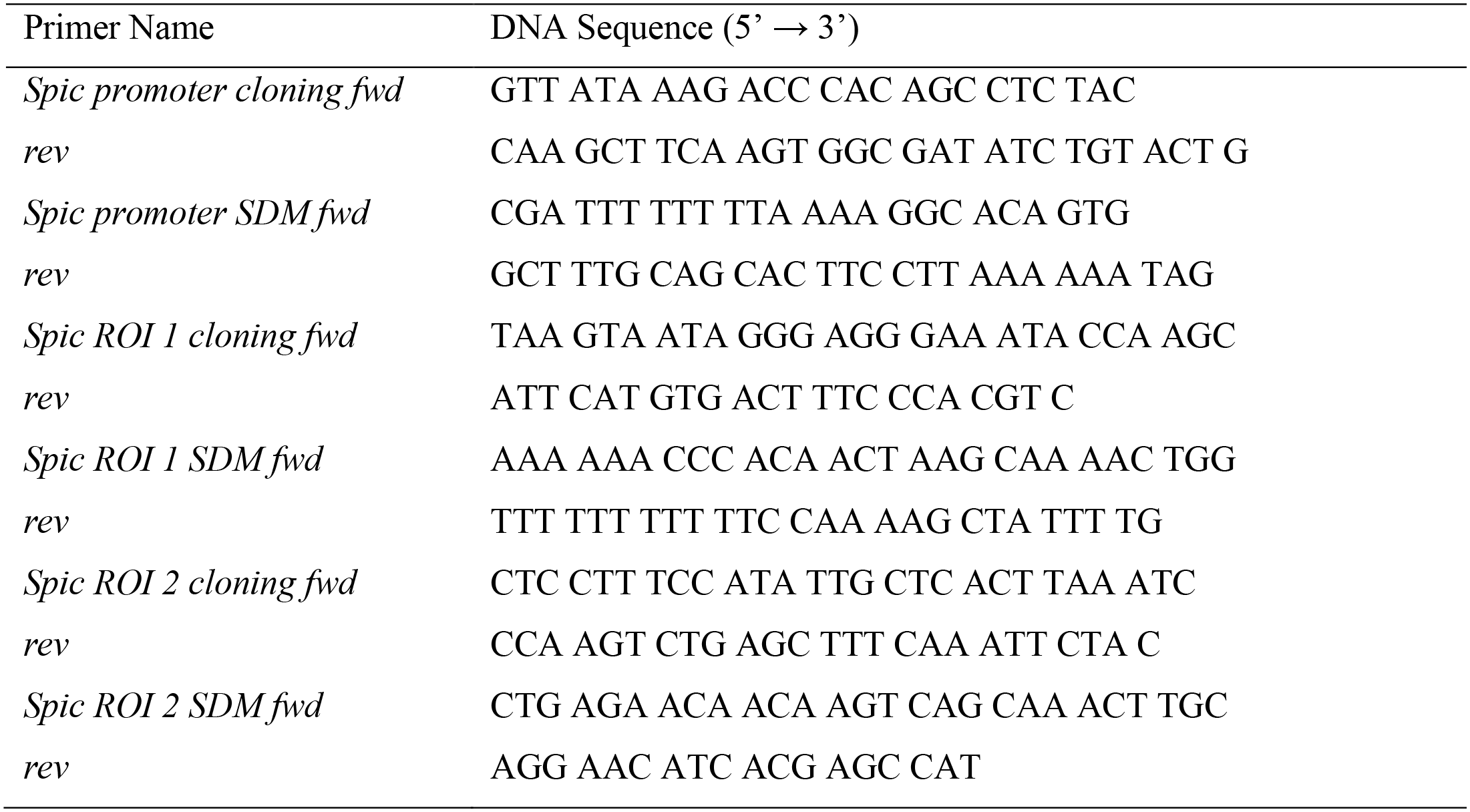
Primer sequences for cloning and site-directed mutagenesis.

**Supplemental Table 3.**
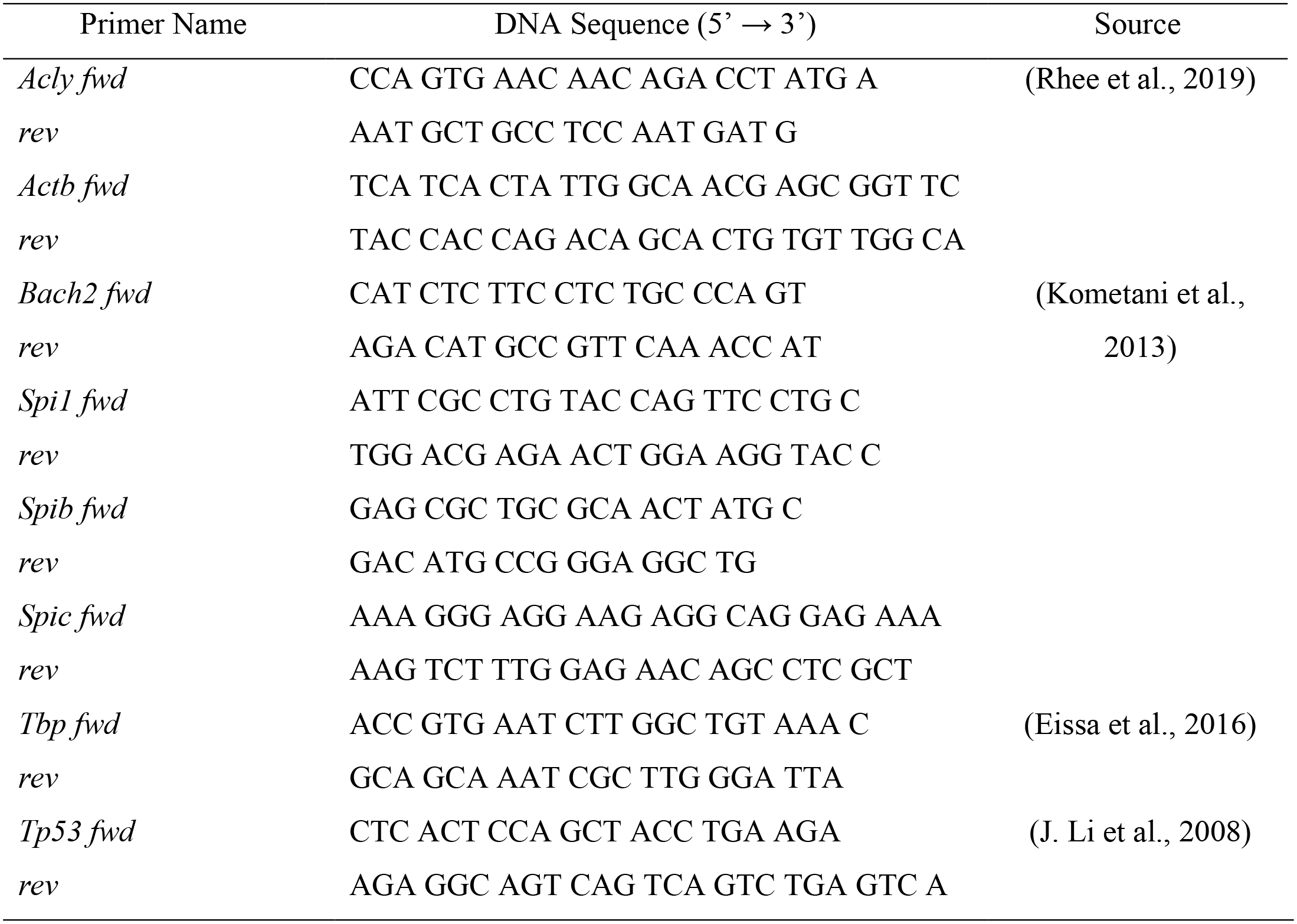
Primer sequences for RT-qPCR.

**Supplemental Figure 1.**
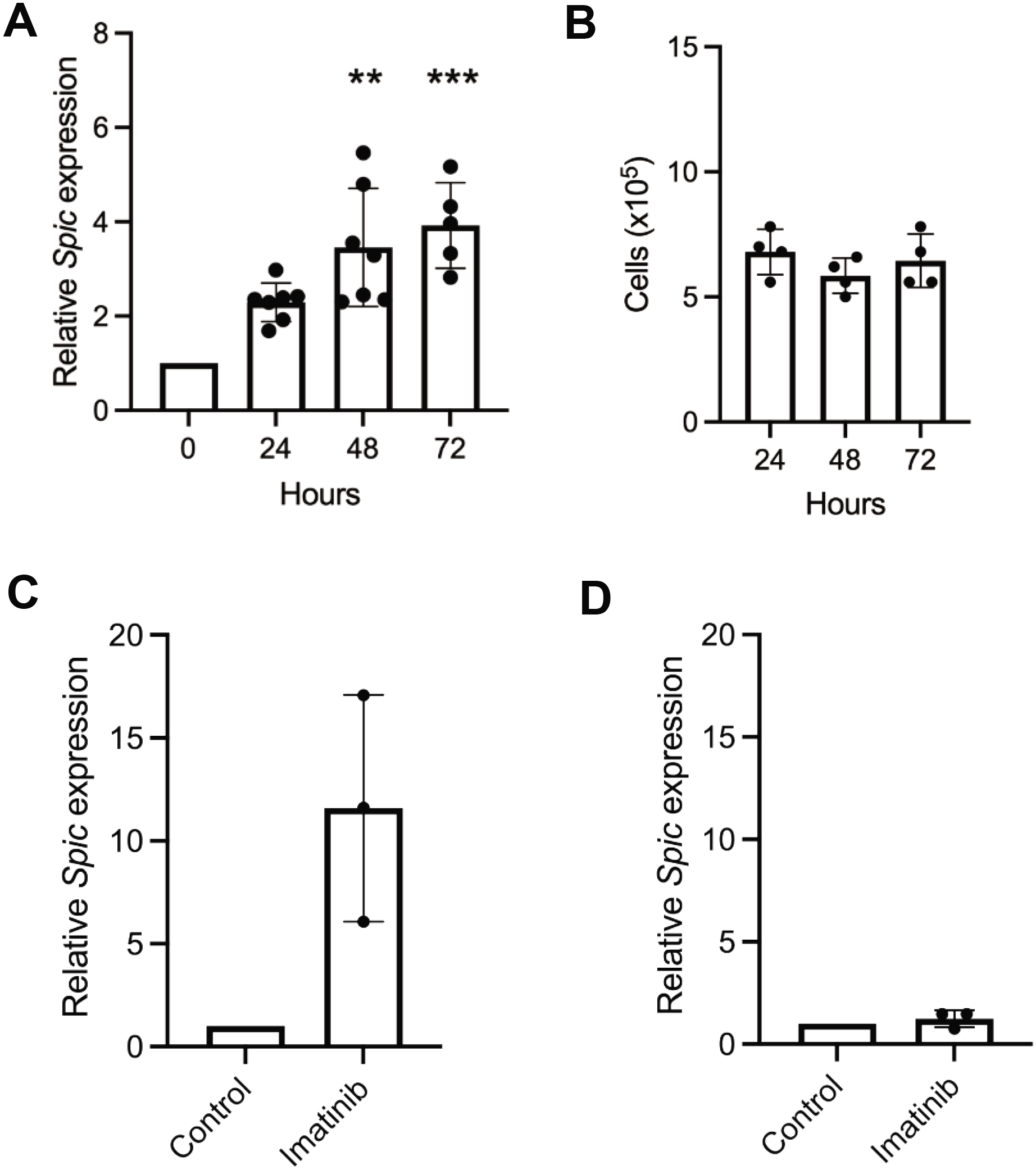
B cells treated with drugs that impair proliferation activate *Spic* expression. (**A**) RT-qPCR analysis of *Spic* expression in WT primary splenic B cells cultured with cytochalasin D for the indicated times. Expression data is relative to freshly isolated B cells and relative to *Tbp*. Data is shown as mean ± SD. Significance was determined using Kruskall-Wallis with Dunn’s multiple comparisons test. (**B**) Live cell counts for cells in (A). Bars indicate mean ± SD. Significance was determined using one-way ANOVA with Tukey’s multiple comparison test. (**C**) RT-qPCR analysis of *Spic* expression in v-Abl-transformed 38B9 pro-B cells or (**D**) IL-7-withdrawn fetal liver-derived WT pro-B cells treated with Imatinib for 24 hours. Data represents mean ± SD. Significance was determined using one sample Wilcoxon test. Expression data for C-D is relative to untreated cells on Day 0 and normalized to *Actb* expression. Individual data points for qPCR experiments represent mean of duplicate wells for one mouse. Cell count data points indicate mean of triplicate counts for each mouse. \*\**p* < 0.01, \*\*\**p* < 0.001.

## Notes

### Competing Interest Statement

The authors have declared no competing interest.

